# Metagenomic Profiling of Bacterial (16S) and Fungal (ITS) Communities on D’Anjou Pears during Long-Term Controlled Atmosphere Storage

**DOI:** 10.1101/2025.11.26.690795

**Authors:** Rawane Raad, Amy Mann, Amrit Pal, Angela Parra, Laura Strawn, Alexis Hamilton, Faith Critzer, Henk C. den Bakker

## Abstract

D’Anjou pears are routinely stored for up to nine months under controlled atmosphere (CA) conditions to meet market demands. While this practice maintains fruit quality, limited information exists on pears’ natural microbiota throughout storage. The objective of this study was to describe fungal and bacterial composition on marketable and unmarketable conventional, whole, intact pears under two storage practices (bulk vs wrapped) at 3, 6, and 9 months in long-term CA cold storage. Storage practices had a significant effect on the composition and succession of both fungal and bacterial communities. Overall, fungal communities exhibited lower estimated Chao1 alpha diversity (mean 18.3) compared to bacterial communities (mean 166.4). No significant differences in Chao1 index were found between the bacterial and fungal communities on marketable or unmarketable pears. Trends in Chao1 indices of fungal and bacterial communities peaked at mid-storage and declined by 9 months, with wrapped pears showing parallel trends and bulk pears exhibiting a sharper late-stage reduction. No distinct clusters could be found for 3- and 6-month fungal communities, irrespective of marketability or being bulk or wrapped. The principal coordinate analysis of the bacterial communities showed tight clustering by time point for the individually wrapped pears, irrespective of their marketability. Bacterial communities included genera common in food-processing and plant environments, such as *Pseudomonas* and *Acinetobacter*. Fungal communities shifted over time, with spoilage-associated genera like *Aureobasidium*, *Penicillium*, *Botrytis*, and *Mucor* present at different storage stages.

**Significance:** This study highlights the influence of storage duration and packaging on microbial succession, establishing initial benchmarks of pear surface microbiomes. The observed lack of significant differences in microbial diversity between marketable and unmarketable pears suggests that these baseline community profiles can serve as critical reference points for identifying other influential factors. Variables such as handling practices may exert a more direct effect on microbial dynamics and, consequently, product quality. Establishing these baselines is essential because they provide a foundation for detecting deviations linked to spoilage or safety risks. Moreover, understanding these patterns can guide the development of targeted microbial control strategies in postharvest systems, enabling interventions that maintain fruit quality, reduce losses, and improve food safety throughout the supply chain.

## INTRODUCTION

Pear (*Pyrus* spp.) production in the United States contributes to 84% of the national fresh pear crop consumption (1). Production is largely concentrated in the Pacific Northwest (Washington State, Oregon, and California) where the industry generates over $250 million in annual sales (1). The 3 main cultivars grown in this region are d’Anjou, Bartlett, and Bosc, with d’Anjou pears representing the largest acreage and highest economic return for U.S. growers (1, 2). To meet year-round market demands and ensure consistent supply, pears must be stored for an extended period of time after harvest. Long-term storage allows growers and distributors to manage seasonal fluctuations in production, maintain fruit availability in off-season markets, and reduce postharvest losses. Controlled atmosphere (CA) storage, which regulates oxygen, carbon dioxide, and temperature, has become the industry standard for preserving fruit quality over several months. Effective long-term storage is a critical component of commercial pear production. It does not only maintain firmness, flavor, and nutritional value but also minimizes economic losses associated with spoilage and decay. Common practices throughout storage include storing the pears in bulk in plastic bins or wrapping each pear with tissue paper impregnated with copper carbonate, food-grade oil, and/or antioxidants such as ethoxyquin to prevent postharvest disorders and the spread of fungal decay organisms (3, 4). These different conditions may play a key role in the microbiota that are present on pear surfaces and how they may change throughout storage duration.

Fruit surfaces host diverse bacterial and fungal communities, collectively known as the epiphytes (5). These microbial communities play important roles in postharvest quality, influencing fruit spoilage, shelf-life, and, in some cases, potential food safety risks. Although no major foodborne outbreaks have been directly associated with pears, other tree fruits such as apples (6) and peaches (7) have been implicated in contamination events, underscoring the need to understand and monitor the microbial populations on pear surfaces. Research on other fruit commodities has increasingly focused on characterizing the complex microbial communities associated with fresh produce and postharvest environments, as these microbiomes influence product quality, safety, and shelf-life. High-throughput sequencing approaches have revealed that fruit surfaces harbor diverse bacterial and fungal taxa beyond what traditional culturing methods can detect, emphasizing the need to consider both culturable fractions and the total microbiota when assessing microbial ecology understanding the fruit microbiome opens new avenues for postharvest biocontrol strategies, shifting the paradigm from targeting single pathogens to managing entire microbial consortia (5). For example, Abdelfattah et al. (8) demonstrated that common postharvest practices such as washing, waxing, and cold storage significantly reshape the apple microbiome, with implications for pathogen suppression and quality maintenance. Al Riachy et al. (9) further showed that long-term storage alters the epiphytic microbiome of apples and influences *Penicillium expansum* occurrence and patulin accumulation, underscoring the dynamic nature of these communities under commercial conditions. Collectively, these findings inform industry practices by suggesting that interventions should account for microbiome shifts over time and under different storage regimes.

Historically, studies of produce-associated microorganisms relied on culture-based methods, which remain the regulatory gold standard for detecting viable pathogens and allow functional characterization of isolates (10). However, these approaches are inherently limited by selective growth conditions and often underestimate microbial diversity, as many taxa are unculturable or require specific growth factors not replicated in vitro (11, 12). In contrast, sequencing-based techniques such as 16S rRNA and ITS amplicon sequencing have revolutionized our understanding of the total microbiota, revealing complex communities that include both dominant and rare taxa (8, 9, 13). Culture-based studies still dominate regulatory testing and targeted pathogen isolation, but sequencing approaches have become the primary tool for hypothesis generation and ecological assessment. This trend underscores a broader movement in food microbiology toward multi-modal strategies that combine culturing and molecular tools to design interventions that maintain fruit quality and safety throughout the supply chain.

Despite the widespread use of CA storage in the pear industry, there is limited information on how such conditions affect the surface microbiome of pears over extended periods. Understanding these potential successional changes in both bacterial and fungal communities is critical for predicting how the microbiome evolves during storage and how it may impact fruit quality. Advances in molecular techniques, including 16S rRNA gene sequencing for bacteria and ITS sequencing for fungi, allow the characterization of these communities (14, 15). Compared to traditional culture-based methods, these approaches capture slow-growing, stressed, or otherwise unculturable taxa, providing a more complete view of the pear surface. Therefore, the objective of this study was to describe fungal and bacterial composition on marketable and unmarketable conventional, whole, intact pears under two different storage practices at 3, 6, and 9 months in long-term CA cold storage to (i) explore the taxonomic composition of the pear surface microbiome and (ii) track community composition of the pear surface microbiome over time. Estimated alpha diversity of ITS1 amplicon sequencing variants (ASVs) for fungal communities was notably lower than that of bacterial communities. The diversity observed in the culturable communities was lower than total rinse communities for both fungi (ITS1) and bacteria (16S). No significant differences were found between the bacterial and fungal communities on marketable or unmarketable pears. Alpha diversity of fungal and bacterial communities peaked at mid-storage and declined by 9 months, with wrapped pears showing parallel trends while bulk pears exhibited a sharper late-stage reduction. This work provides the primary baseline data on pear fungal and bacterial diversity and composition, which can be foundational in the development of effective microbial management strategies. From an industry perspective, understanding the temporal shifts of the pear surface microbiome can inform storage practices, microbial risk assessment, and potential biocontrol strategies by identifying critical windows of community change, predict conditions that favor pathogen establishment, and guide interventions that promote beneficial taxa while suppressing spoilage organisms. Such insights enable the development of targeted treatments, optimized storage parameters, and microbiome-informed quality control protocols that reduce postharvest losses and enhance food safety.

## RESULTS

### Alpha diversity

Estimated Chao1 alpha diversity of ITS1 amplicon sequencing variants (ASVs) for fungal communities was notably lower (average 18.3, CI 15.5 – 21.1) than estimated alpha diversity of 16S ASVs in bacterial communities (average 166.4, CI 136.2 – 196.6). The diversity observed in the culturable communities was significantly (*P* < 0.01) lower than total rinse communities for both fungi (ITS1) and bacteria (16S). For example, total bacterial communities showed an average of 298 ASVs, while bacterial communities recovered from cultures showed an average of 32 ASVs. The average number of fungal (ITS1) ASVs was 25 for DNA isolated from the total rinse, and 6 for DNA isolated from cultures. No significant differences in alpha diversity of fungal communities were found between bulk and individually wrapped pears. Bacterial communities did show a significant difference (*P* < 0.05) with bulk (182.51 ASVs) having a higher average diversity than wrapped (105 ASVs). No significant differences in Chao1 richness indices were observed when marketable versus unmarketable pears were compared for both fungal and bacterial communities.

Using the total rinse richness estimates, we examined for significant differences in richness between the 3-, 6-, and 9-mo time points. Bulk and wrapped pears were tested separately. For the fungal communities in bulk pears, we saw no significant change in the Chao1 index between the 3- and 6-mo time points, while a significant (*P* < 0.05) decrease was observed from the 6 to 9-mo time points. Wrapped pears showed a different pattern; a significant increase in ASV richness was observed between the 3- and 6-mo time points, followed by a significant decrease between the 6- and 9- mo time points. For bacterial communities in bulk pears, a significant (*P* < 0.05) decrease in richness of ASVs was observed between the 3- and 6-mo time points, while a significant (*P* < 0.05) increase was observed between the 6- and 9- mo time points. For bacterial communities on wrapped pears, a similar pattern to the observed fungal communities on wrapped pears was observed, a significant increase in richness between the 3- and 6- mo time points, and a significant decrease between the 6- and 9- mo time points.

### Beta diversity

No distinct clusters could be found for 3- and 6-mo fungal communities (Fig. 1), irrespective of marketability or storage condition (bulk or wrapped). Fungal communities of 9-mo marketable bulk pears and unmarketable wrapped pears are distinct from the 3- and 6-mo pears (Fig. 1). A Constrained ordination (CAPscale) analyses based on both bulk and wrapped pears showed that marketability and pears being individually wrapped had a highly significant effect on the beta diversity (*P* < 0.01), while time of sampling had a significant effect at *P* = 0.045. When individually wrapped pears and bulk pears were analyzed separately, time of sampling was still highly significant. Marketability was slightly less significant for bulk pears (*P* = 0.05), while it was not significant for individually wrapped pears (*P* = 0.65).

**Fig. 1:**
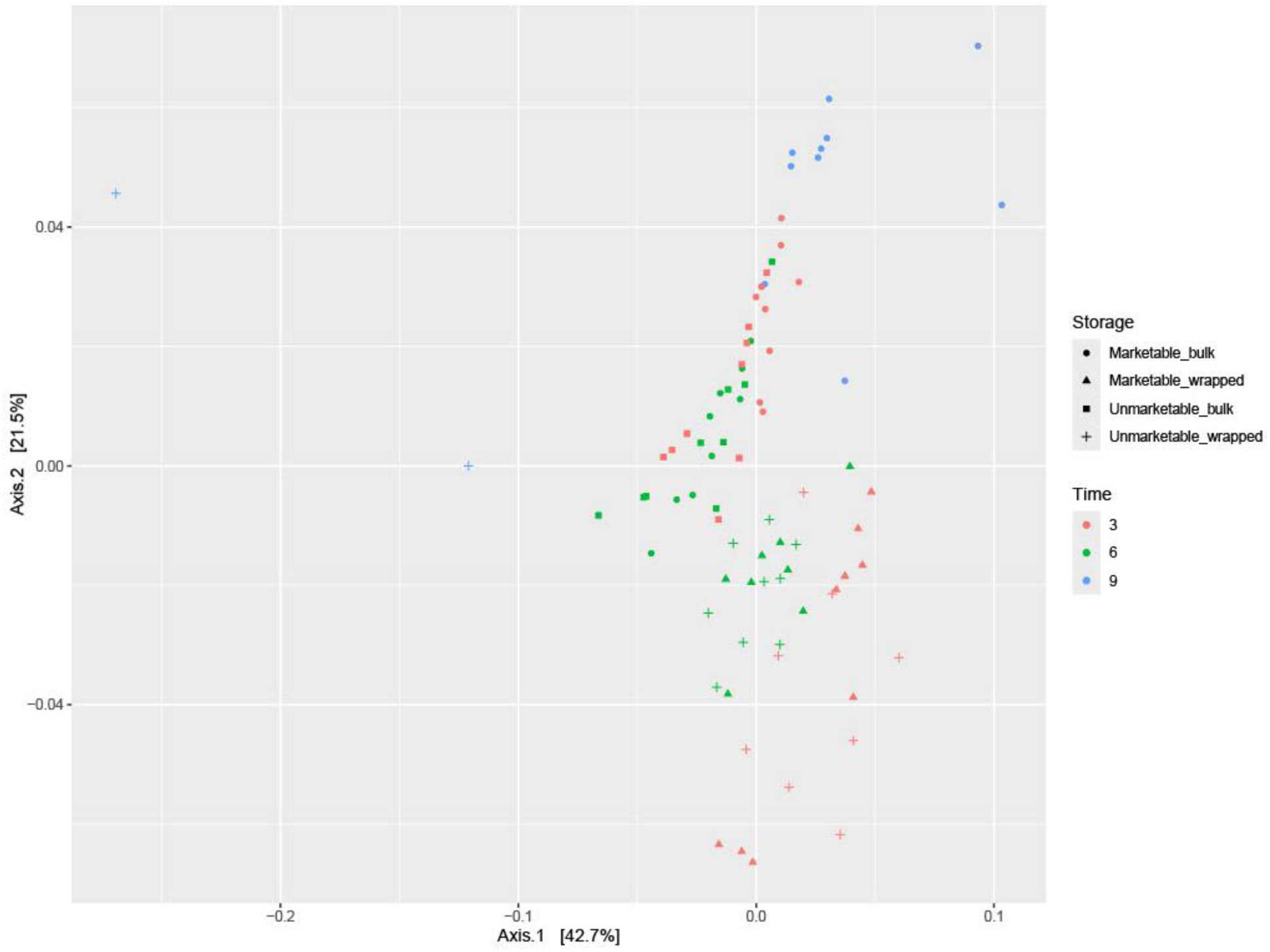
Principal Coordinate Analysis (PCoA) of fungal (ITS) beta diversity (UniFrac) from pear rinse samples, grouped by storage time (3-, 6-, and 9-months time period), marketability, and packaging condition (wrapped vs bulk)

Principal coordinate analysis (PCoA) of the bacterial communities showed tight clustering by time point for the individually wrapped pears, regardless of marketability (Fig. 2). The 9-mo marketable bulk pears cluster with the 9-mo individually wrapped pears, while 3- and 6-mo bulk pears form very loose clusters which are mainly separated on the Y-axis, with no clear clustering by marketability (Fig. 2). CAPscale analyses confirm the same pattern, showing that both time and being either individually wrapped or bulk have a highly significant (*P* < 0.01) effect on the beta diversity of samples, while marketability is not significant (*P* > 0.05).

**Fig. 2:**
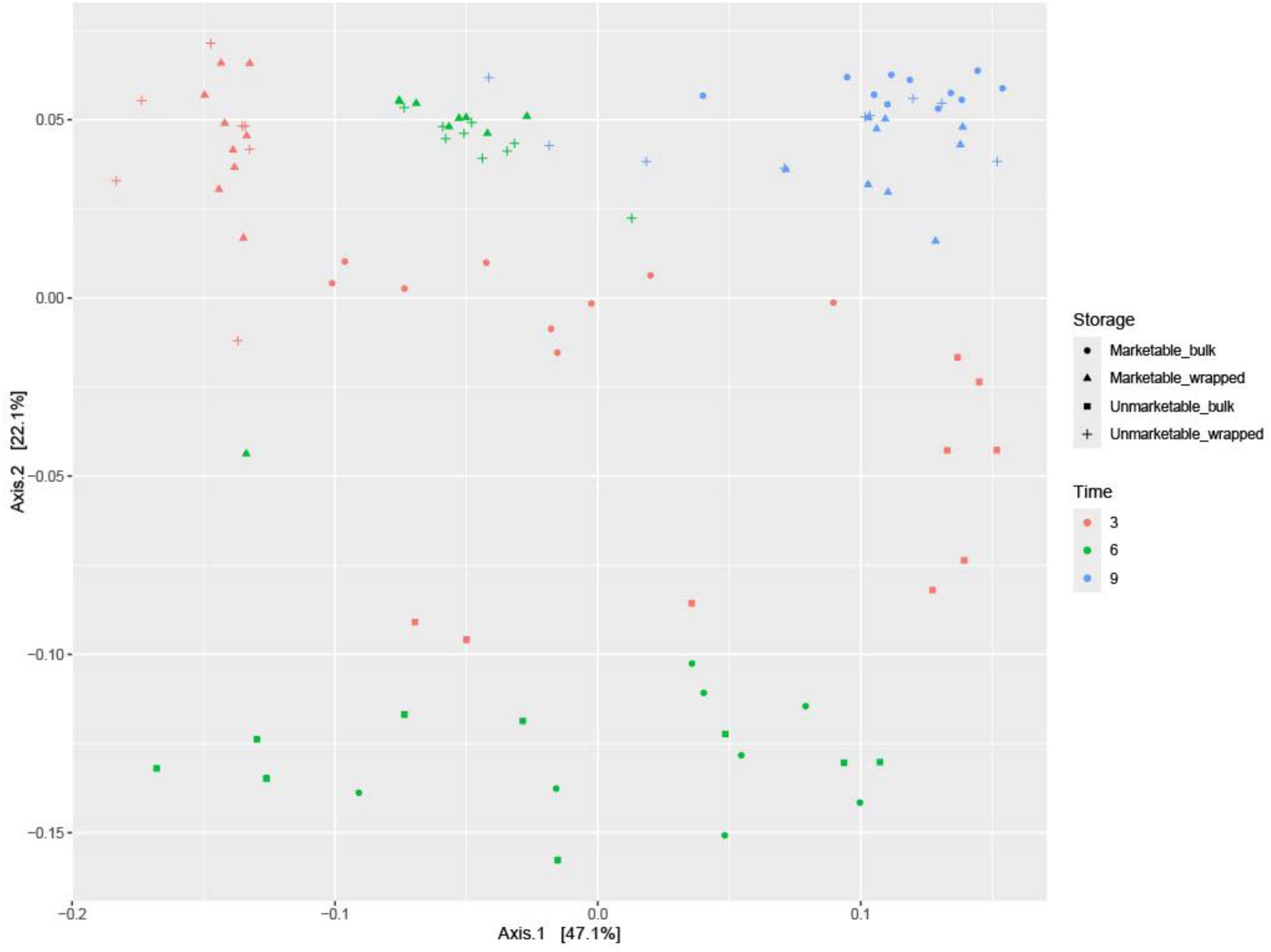
Principal Coordinate Analysis (PCoA) of bacterial (16S) beta diversity (UniFrac) from pear rinse samples, grouped by storage time (3-, 6-, and 9-months time period), marketability, and packaging condition (wrapped vs bulk)

### Composition of fungal and bacterial communities associated with the surface of pears

Bacterial communities (Fig 3.) consisted of a mixture of genera found in food processing environments and other build environments such as *Acinetobacter*, *Pseudomonas*, bacteria associated with the microbiota of plants (e.g., *Frondihabitans*, *Erwinia*) and sugar rich environments (e.g., *Gluconobacter*). The fungal communities (Fig. 4) were dominated at the 3-and 6-mo time point by *Aurobasidium* and *Penicillium* species, however these genera were virtually absent at the 9-mo time point. At all time points we found a large variety of psychrophilic and psychrotrophic yeasts (e.g., *Mrakia*, *Leucosporidium*, *Cutaneotrichosporon*, *Tausonia*). Additionally fungal genera associated with fruit spoilage were found, such as *Botrytis* and *Mucor*. For bulk pears ASVs associated with the bacterial genera *Gluconobacter*, *Rhanella*, *Leuconostoc* and *Pantoea* were significantly enriched in 6-mo communities compared to 3-mo communities, while *Pseudomonas* were significantly reduced. In the 9- versus 6-mo comparison ASVs associated with *Pseudomonas*, *Acinetobacter*, *Microbacter* and *Rhodococcus* were significantly enriched, while *Gluconobacter* and *Rhanella* were reduced. For fungal communities in bulk pears ASVs associated with *Neonectria*, *Niesslia*, *Penicilllium* and *Fusarium* were significantly enriched in 6-mo communities as compared to 3-mo communities, while ASVs associated with *Penicilllium*, *Aureobasidium, Mrakia*, *Botrytis* and *Candida* were significantly reduced. ASVs associated with *Leucosporidium* were significantly enriched in 9-mo fungal communities compared to 6-mo communities, while ASVs associated with *Aureobasidium*, *Penicillium fuscoglaucum* and *Neonectria* were significantly reduced.

**Fig. 3:**
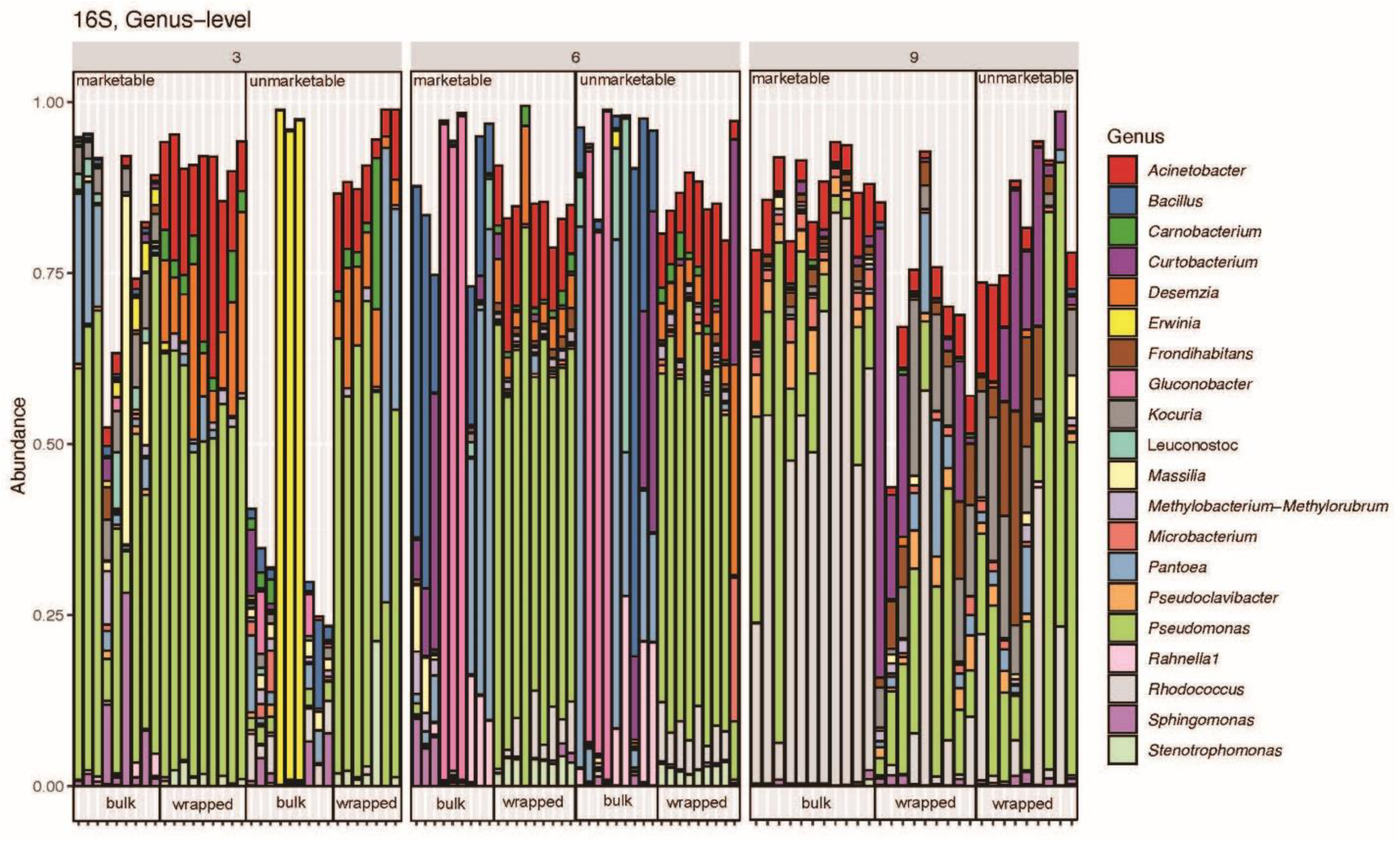
Genus-level composition of bacterial communities on pear rinse surfaces across marketable and unmarketable fruit, stored in bulk or wrapped conditions over 3-, 6-, and 9-month periods

**Fig. 4:**
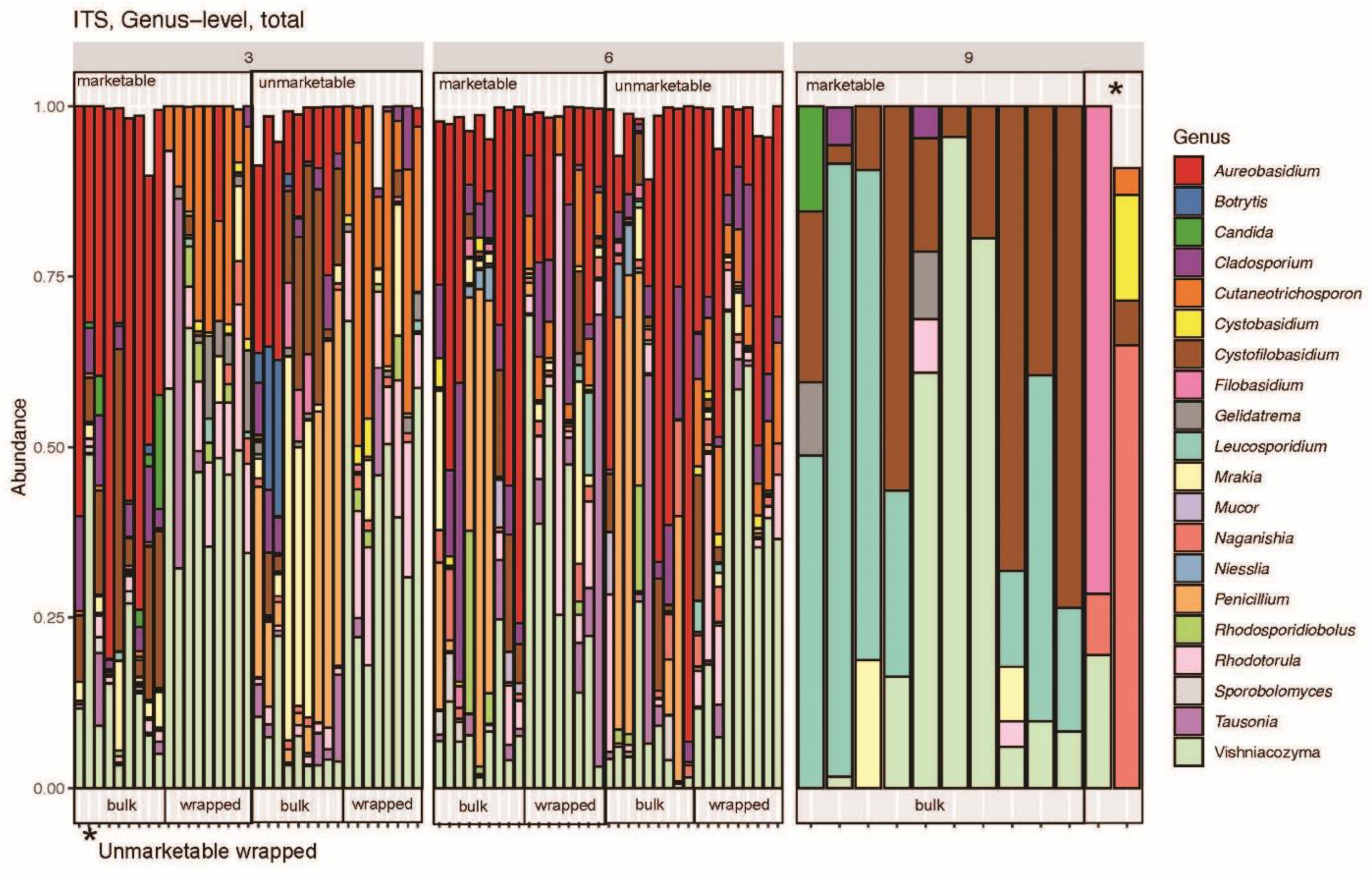
Genus-level composition of bacterial communities on pear surfaces across total rinses of marketable and unmarketable fruit, stored in bulk or wrapped conditions over 3-, 6-, and 9-month periods

For individually wrapped pears ASVs associated with the genera *Williamsia*, *Pseudomonas*, *Frondihabitans*, *Rhodococcus* and *Curtobacterium* were significantly enriched in 6-mo bacterial communities, while ASVs associated with *Pseudomonas*, *Stenotrophomonas* and *Carnobacterium* were reduced. In the 9-mo versus 6-mo comparison ASVs associated with *Pseudomonas*, *Acinetobacterium*, *Microbacterium* and *Rhodococcus*, while *Rhanella* and *Gluconobacter* associated ASVs were significantly reduced. In fungal communities in individually wrapped pears ASVs associated with *Cadophora*, *Mrakia*, *Cladosporium* and *Vishniacozyma* were significantly enriched in 6-mo communities, and only one ASV associated with the yeast genus *Metschnikowia* was significantly reduced. In 9-mo communities compared to 6-mo communities, only one ASV associated with the genus *Filobasidium* was enriched, while ASVs associated with *Aureobasidium*, *Vishniacozyma*, *Cutaneotrichosporon*, and *Rhodotorula* were among the genera associated with reduced ASVs.

## DISCUSSION

D’Anjou pears are routinely stored for up to 9 months under CA conditions to meet market demands. Limited information exists on pear’s natural microbiota throughout storage. Fungi are important contributors to pear quality and bacteria can influence not only quality but also the safety of pears. Microbial taxa are commonly characterized using ASVs (16–19). ASVs are often used to estimate species richness, which can then be compared across samples from the environment (e.g. fruits, vegetables, water, or soil, etc.). The Chao1 index is a richness estimator that accounts for unseen or rare taxa by incorporating the number of singletons and doubletons (20). In this study, the estimated Chao1 diversity of ITS1 ASVs for fungal communities was notably lower than the estimated diversity of 16S ASVs in bacterial communities. The microbial load on fruits can vary widely depending on the type of fruit, its surface characteristics, growing conditions, and handling (18, 19, 21). Fungal colonization of fruit surfaces tends to be more selective than bacterial colonization (19, 22, 23). Fungal growth on pears, particularly during prolonged storage, is often dominated by a few genera that are well adapted to the fruit environment, such as *Penicillium*, *Botrytis*, or *Alternaria* (24). These fungi can outcompete other taxa through rapid sporulation, enzymatic degradation of fruit tissues, or production of antimicrobial metabolites (25–28), resulting in reduced overall fungal richness. In this study, a large variety of psychrophilic and psychrotrophic yeasts (e.g., *Mrakia*, *Leucosporidium*, *Cutaneotrichosporon*, and *Tausonia*) were observed. *Aurobasidium* and *Penicillium* species were most dominant at 3- and 6-mo time points (Fig. 2). *Aureobasidium* is a ubiquitous yeast-like fungus often found on fruit surfaces, known for biofilm formation and tolerance to stress, contributing to long-term survival (29, 30). *Penicillium* is a filamentous fungus responsible for blue and green mold rots on pears and apples, producing enzymes and mycotoxins that drive decay (31, 32). *Botrytis* and *Mucor* were also found (Fig. 2), both spoilage molds that cause gray and soft rot, respectively (33–35). By contrast, bacterial communities on fruit surfaces are typically more diverse, comprising both epiphytic and environmental species, many of which can persist under CA storage without necessarily dominating the community. In this study, the composition of bacterial communities associated with the surface of pears consisted of a mixture of genera found in food processing environments and other built environments, such as *Acinetobacter*, a genus of Gram-negative bacteria commonly found on plant surfaces and in soil (36). *Pseudomonas*, a diverse group of Gram-negative bacteria frequently isolated from fruits and vegetables, with some species contributing to spoilage through enzymatic degradation and biofilm formation (37, 38) were also detected. Other bacteria associated with the microbiota of plants (e.g., *Frondihabitans*, *Erwinia*) and sugar-rich environments (e.g., *Gluconobacter*) were also found. Reduced oxygen and elevated carbon dioxide are effective at suppressing fungal growth, particularly of aerobic saprophytic fungi such as *Penicillium*, *Botrytis*, and *Mucor* (39–41). Storage conditions may differentially influence fungi and bacteria. This difference underscores the need for targeted strategies: fungal control may require interventions aimed at a few dominant pathogens, while bacterial management must consider a broader, more resilient community. Understanding these dynamics is critical for designing microbiome-informed approaches to improve pear quality and reduce postharvest losses.

Reduced oxygen storage is widely recognized as an effective strategy to delay fungal growth and extend fruit shelf life. CA systems typically maintain oxygen levels between 1–3%, which significantly inhibits pathogens such as *Penicillium digitatum* and *P. italicum* without compromising fruit quality (42, 43). For example, Archer et al. (42)demonstrated that oranges stored under low oxygen (0.9 kPa O₂, ∼ 0.9%) exhibited markedly reduced mold lesion development compared to untreated controls, while key quality attributes remained largely unaffected. Similarly, combining CA storage with biocontrol agents such as antagonistic yeasts has proven effective in apples and pears, where 3% O₂ and elevated CO₂ levels suppressed gray mold and blue mold and enhanced yeast efficacy (43) These findings underscore the importance of oxygen management as a non-chemical intervention for postharvest disease control. By contrast, Hoogerwerf et al. (2020) reported an atypical modified atmosphere containing 80% oxygen and 20% CO₂, which inhibited *Botrytis cinerea*, *Penicillium discolor*, and *Rhizopus stolonifer* for up to 17 days, while elevated CO₂ alone suppressed growth for about 11 days. This high-oxygen approach differs fundamentally from conventional CA strategies, which rely on hypoxic conditions, and highlights the diversity of atmospheric manipulations being explored for decay control. However, such treatments may also influence bacterial taxa differently, as genera like *Pseudomonas* and *Acinetobacter* can adapt to modified atmospheres through metabolic flexibility and biofilm formation (44–46). This suppression of fungi under CA atmosphere storage could therefore have limited the number of detectable fungal ASVs compared to bacteria.

When comparing communities obtained from culturable samples with those detected in whole-pear rinses, alpha diversity indices consistently showed fewer total ASVs in the culturable fraction. This highlights the value of whole-fruit rinsing approaches and underscores the biases inherent to culture-based methods, which often favor fast-growing or easily culturable taxa prioritized by target growth conditions (e.g., *Pseudomonas*, *Enterobacteriaceae*) while underrepresenting slow-growing, stressed, or obligate symbiotic organisms (11, 12, 47, 48). For example, Ruiz Rodriguez et al. (48) used both culture-dependent and shotgun metagenomic (culture-independent) approaches to characterize microbial communities on tropical fruit surfaces (e.g., guava, papaya, medlar flowers). They found that while the most abundant species were identified by both methods, less abundant taxa were often missed by culture-based methods, likely due to selective enrichment in cultures. Total rinse approach will avoid a direct bias imposed by cultivation media used in a culture-dependent (enrichment) strategy and should be the method utilized in future studies when the objective is to characterize community dynamics irrespective of target organisms or indicators (e.g., generic *Escherichia coli*).

Storage placement conditions such as bulk vs individually wrapped had no effect on fungi diversity over time, but bacterial communities showed a significant difference (*P < 0.05)*. Wrapping can create microenvironments with altered humidity and gas exchange (49), potentially selecting for specific fungal taxa while excluding others. In bulk storage, fungal spores may disperse more easily but remain dominated by a few highly competitive taxa, leading to relatively low richness. The decline in fungal alpha diversity observed in bulk-stored pears between 6 and 9-mo can be explained by both physiological constraints of the storage environment and competitive ecological dynamics among fungal taxa. Prolonged exposure to low O₂ and elevated CO₂ conditions suppresses the growth of many aerobic organisms, creating a selective environment where a subset of storage-adapted fungi, such as *Penicillium*, *Alternaria*, and *Botrytis*, can persist and eventually dominate, as discussed previously (25, 31–34). Such selective pressure likely explains the significant reduction in community richness as storage time progresses, particularly after 6 months. Taxon-specific changes over time also support this interpretation. At 6 months, ASVs affiliated with *Neonectria*, *Niesslia*, and *Fusarium* were significantly enriched compared to 3 months, suggesting these fungi are particularly well-adapted to intermediate storage conditions.

Bacteria can more readily occupy micro-niches across fruit surfaces and persist in a broader range of storage conditions, maintaining higher observed diversity. In the fresh pear industry, wrapping papers impregnated with copper carbonate, antioxidants (e.g., ethoxyquin), and oils have been widely used for over a century to reduce physiological disorders such as superficial scald (3, 50). However, assessment of wrapping paper efficacy on fresh pears have focused exclusively on *B. cinerea* (51), leaving broader microbial impacts largely unexplored. In this study, both wrapped and bulk pears were stored under CA conditions which are known to suppress many epiphytic taxa, including genera such as *Sphingomonas* and *Methylobacterium*. The early decline in alpha diversity observed in bulk pears between 3- and 6- mo may therefore reflect the combined effects of CA conditions and nutrient limitation, rather than packaging alone. This distinction is important because while wrapping papers may influence surface chemistry and microenvironment, the overarching atmospheric conditions likely exert a stronger selective pressure on microbial communities. As tissues soften/sense, micro-leaks of sugars/phenolics create new niches (52). This can enable secondary colonizers and metabolically flexible taxa (e.g., *Pseudomonas*, *Pantoea/Enterobacteriaceae*, *Acinetobacter*) to rebound, raising richness. Similar patterns have been documented in pears treated with 1-methylocyclopropene, a synthetic plant growth regulator. The study reported that fungal pathogens such as *Botryosphaeria*, *Penicillium*, and *Fusarium* were suppressed compared to untreated fruit (19). Apples also show strong temporal succession in storage, with stable bacterial (*Pseudomonas*, *Pantoea*) and fungal (*Aureobasidium*, *Cladosporium*) cores persisting while other taxa turn over across cold storage (53, 54). Together, these findings highlight that fruit microbiomes are strongly shaped by storage environment, and that packaging type can influence its corresponding microbial community. Future work should pair diversity analyses with metabolite profiling to link taxonomic shifts to function and track the relative abundance of sentinel genera such as *Pseudomonas*. Additionally, examining biofilm markers (e.g., extra-polymeric substance genes, microscopy) would help determine whether late-stage dominance coincides with biofilm formation, particularly in wrapped fruit.

Marketability of fruits typically aligns with standards used to assess consumer acceptability, which are based on visual attributes and not on food safety considerations. This includes blemishes, patches of rot or mold, sun- or frost-burn, bruising, or scarring, etc. The absence of significant differences in alpha diversity between marketable and unmarketable pears suggests that the transition from marketable to unmarketable may not depend on the total diversity of surface-associated microbes, but rather on the activity and behavior of specific taxa. For example, unmarketable pears may reflect microbial processes such as the internalization of microbes and pathogens into fruit tissues (55, 56) or the establishment of biofilms, which would not necessarily be captured by surface-level diversity measures.

The fungal communities associated with pears showed dynamic changes over storage, but distinct clustering patterns were only observed at the 9-mo time point (Fig. 1). Specifically, fungal communities from 9-mo marketable bulk pears and unmarketable wrapped pears were clearly separated from those at 3- and 6-mo, irrespective of marketability or packaging. CAPscale analyses revealed that both marketability and packaging type (bulk vs. individually wrapped) had a highly significant effect on fungal beta diversity (*P < 0.01*), while time of sampling was less significant (*P = 0.045*). When pears were analyzed by packaging type, time remained a strong driver of fungal community structure, while marketability was less significant for bulk pears (*P = 0.05*) and not significant for wrapped pears (*P = 0.65*). These results suggest that packaging and marketability jointly influence fungal community composition, with marketability exerting a stronger effect in bulk fruit, likely due to greater exposure or handling differences, whereas time consistently shaped communities across treatments.

For bacterial communities, ordination analyses demonstrated tighter clustering by storage time for individually wrapped pears, regardless of marketability, indicating that temporal dynamics outweighed quality status. By contrast, bulk pears at 3- and 6-mo exhibited loose clustering, with no clear separation by marketability, whereas 9-mo marketable bulk pears clustered closely with 9-mo wrapped pears. CAPscale results confirmed these patterns: both time and packaging type significantly influenced bacterial beta diversity (*P < 0.01*), while marketability showed no significant effect (*P > 0.05*). Together, these findings indicate that while time and packaging are consistent drivers of microbial succession in stored pears, marketability is more relevant for fungal than bacterial communities, particularly in bulk fruit.

This study provides a valuable baseline for understanding pear-associated microbial communities using a microbiome approach, characterizing these communities as they naturally exist without targeted plant or human pathogens. Such baseline data are critical for future efforts to link microbial ecology with fruit physiology, spoilage dynamics, and food safety outcomes. However, several limitations must be acknowledged. Sampling was opportunistic and involved multiple pear lots rather than tracking a single batch longitudinally, limiting the ability to infer precise temporal dynamics. Although pears from the same variety and storage conditions are expected to behave similarly, batch-level variability cannot be fully excluded. Additionally, amplicon sequencing captures only a subset of the microbial community, potentially overlooking functionally important but low-abundance taxa or those with atypical marker genes. This limitation is common in microbiome studies and highlights the need for complementary approaches such as shotgun metagenomics (13). Another important consideration is that while diversity and community composition were characterized, functional implications, such as metabolite shifts, pathogen suppression, or interactions with fruit physiology, were not directly tested. Microbiome data alone cannot predict whether observed taxa contribute to spoilage or confer protective effects against pathogens. Future research should therefore integrate multi-omics approaches, including metabolomics and transcriptomics, to link microbial succession with biochemical changes during storage. Experimental inoculations under controlled conditions could further clarify causal relationships between microbiome composition and outcomes such as decay incidence or patulin accumulation, as demonstrated in other studies (8, 9, 57)

Understanding the temporal shifts of the pear surface microbiome can inform storage practices, microbial risk assessment, and potential biocontrol strategies by identifying critical windows of community change. This information helps to build our foundational knowledge and improve our ability to predict conditions that favor pathogen establishment and guiding interventions that promote beneficial taxa while suppressing spoilage organisms. Such insights could lead to microbiome-informed storage protocols, targeted treatments, and improved quality control measures that reduce postharvest losses and enhance food safety.

## CONCLUSION

In conclusion, no significant differences in Chao1 alpha diversity index were observed in the overall bacterial and fungal community structures between marketable and unmarketable pears, despite the presence of known spoilage fungi such as *Botrytis*, *Mucor*, and *Penicillium*. Instead, storage practices played a more prominent role, with packaging type (individually wrapped vs bulk) significantly shaping the composition and succession of both fungal and bacterial communities. For fungi, alpha diversity peaked at 6 months before declining at 9 months, while bacterial communities from individually wrapped pears followed a similar temporal trend. In contrast, bulk pears only showed a significant decline in bacterial diversity between 6 and 9 months. These findings underscore the critical influence of storage conditions on microbiome dynamics and highlight the need to consider microbial ecology in postharvest management strategies. By providing the first comprehensive baseline of pear-associated microbiomes under commercial storage, this study lays the foundation for future research aimed at linking microbial succession with fruit physiology, spoilage risk, and food safety. Such knowledge is essential for developing microbiome-informed interventions that optimize storage practices, reduce losses, and enhance the quality and safety of pears in the supply chain.

## MATERIALS AND METHODS

### Pears

D’Anjou pears were shipped from an industry collaborator in Washington State to the Center for Food Safety (Griffin, GA) at 3, 6, and 9- mo after harvesting. Pears (N = 256) were either received in bulk or individually wrapped; for both categories pears were marked as ‘marketable’ or ‘unmarketable’, as determined by industry collaborators. Marketability typically aligns with standards used to assess consumer acceptability, which are based on visual attributes and not on food safety considerations. Marketable and unmarketable pears were collected for each time point and type of pear, with the exception of bulk 9-mo pears, for which only marketable fruit was obtained.

### Bacterial and fungal isolation

Five pears per lot of the same marketability type were sonicated with 250 mL of wash solution comprising 1X Tris-EDTA buffer (G-Biosciences, Saint Louis, MO, USA) supplemented with 2% Tween 80 (Research Products International, Mount Prospect, IL, USA) in a stomacher bag. To dislodge microorganisms from the surface of the samples, one individual pear was sonicated per minute for a total of five pears/minutes (VWR International, LLC, Radnor, PA, USA). During sonication, samples were securely contained within sterilized sampling bags to maintain aseptic conditions. To recover the culturable microbiota, 10 mL of the sample was collected, and 4 mL aliquots were plated in duplicate onto Tryptic Soy Agar (TSA; Difco, Becton Dickinson Co, Franklin Lakes, NJ, USA). The TSA plates were incubated at 35°C for 24 hours to allow bacterial growth. For culturable fungi, 1 mL of the same (4 mL) aliquot was plated in duplicate onto Potato Dextrose Agar (PDA; Difco, Becton Dickinson Co), and the plates were incubated at 25 °C for 7 days.

### DNA isolation of microbial communities of the pear surface

The remaining wash solution (∼ 235 mL) was divided into seven aliquots of 30–35 mL in sterilized conical centrifuge tubes. These aliquots were centrifuged at 3180 x g for 15 minutes at 4 °C. Following centrifugation, the clear supernatant was carefully removed, leaving a dense pellet in each aliquot. The pellets from all aliquots were combined into a single tube and centrifuged again at 3180 x *g* for 15 minutes at 4 °C. After discarding the supernatant, the consolidated pellet was transferred to a microcentrifuge tube and subjected to a final centrifugation at 10,000 × *g* for 10 minutes. The remaining concentrated pellet served as the sample for DNA extraction.

For the culturable microbiota, 1 mL of Phosphate Buffered Saline (PBS; Difco, Becton Dickinson Co) was pipetted onto the TSA and 5 mL of PBS onto the PDA plate surface, and the bacterial lawn was gently removed using a sterile L-shaped spreader. The resulting suspension was transferred into 2 mL tubes for each sample. The pooled solution was then concentrated by centrifugation at 10,000 × *g* for 5 minutes at 4 °C and stored at −20 °C until DNA extraction. DNA was extracted from the pellet using the DNeasy PowerSoil Kit (Qiagen, Hilden, Germany) in accordance with the manufacturer’s instructions.

### Amplicon sequencing of bacterial and fungal communities

High throughput sequencing was performed on the Illumina MiSeq sequencer at the University of Georgia’s Center for Food Safety, generating paired 300bp reads for each amplicon. To target bacterial communities the V3-V4 region of 16S rRNA was sequenced, while the first internal transcribed spacer (ITS1) of the eukaryotic ribosomal cistron was targeted for the fungal communities. Primers and PCR parameters for both V3-V4 16S and ITS amplicon sequencing were obtained from Illumina application notes.

### Bioinformatics and statistical analyses

All bioinformatics and statistical analyses, with the exception of sequence alignment and phylogenetic inference were performed in R (58). Sequence alignment was performed using Muscle 5.1 (59) and phylogenetic trees were inferred using FastTree (60) on a Linux server. The R package DADA2 (61) was used to infer amplicon sequence variants largely following the tutorials outlined on the website of the authors (https://benjjneb.github.io/dada2/tutorial_1_8.html). Phyloseq (62) was used for further microbiome analyses, largely following the workflow outlined in Callahan et al. 2016 (63). DeSeq2 analyses for individual time points were performed separately for bulk and individually wrapped pears to infer which ASVs were enriched in the communities at different time points. Overrepresented taxa in pairwise comparisons (e.g., month 3 versus 6, marketable versus unmarketable were also inferred using the DeSeq2 R package (64), while CAPscale analyses in the vegan R package (65) were performed to infer which parameters were significantly influencing beta-diversity in the microbial communities.

Alpha diversity of amplicon sequence variants (ASVs) was estimated using the Chao1 statistic The Chao1 index is widely used in microbiome studies, including food systems, because it estimates species richness while accounting for undetected rare taxa, which is critical in environments where sequencing depth may miss low-abundance organisms. This helps researchers avoid underestimating diversity and provides a more accurate baseline for comparing treatments or conditions (16, 66, 67). Alpha diversity indices were compared between two groups (e.g., marketability, packaging type, total rinse vs culturable) using Welch’s t-test, which accounts for unequal variances. To assess changes in alpha diversity over time, ANOVA followed by Tukey’s post hoc test was used. To compare the composition (beta diversity) of the fungal and microbial communities found on the pear surfaces PCoA were performed based on weighted UniFrac (68) distances. To test which parameters (e.g., marketability, bulk versus wrapped) were significantly associated with beta diversity we used a constrained ordination via CAPscale analysis in the vegan package. Beta diversity analyses were only performed for the ‘total rinse’ samples, as the datasets of culturable organisms only represent small subsets of the total diversity. *P < 0.05* was considered significant.

## ACKNOWLEDGMENTS

The authors express their gratitude to industry collaborators from the Pacific Northwest pear industry for their invaluable input that contributed to this study. The authors would also like to acknowledge Dr. Claire Murphy from the University of Washington for her help with organizing pear shipments to the appropriate processing labs.

## Author contributions: CRedit

**Rawane Raad:** Investigation, Writing – original draft preparation, Writing – reviewing and editing. **Amy Mann**: Investigation, Writing – reviewing and editing. **Amrit Pal:** Investigation. **Angela Parra:** Investigation, **Laura Strawn**: Supervision, Project administration, Funding acquisition, Writing – reviewing and editing**. Alexis Hamilton**: Supervision, Writing – reviewing and editing **Faith Critzer:** Supervision, Writing – reviewing and editing. **Henk C. den Bakker**: Conceptualization, Methodology, Software, Formal Analysis, Resources, Supervision, Writing – review & editing

